# Probing the limits of alpha power lateralization as a neural marker of selective attention in middle-aged and older listeners

**DOI:** 10.1101/267989

**Authors:** Sarah Tune, Malte Wöstmann, Jonas Obleser

## Abstract

In recent years, hemispheric lateralization of alpha power has emerged as a neural mechanism thought to underpin spatial attention across sensory modalities. Yet, how healthy aging, beginning in middle adulthood, impacts the modulation of lateralized alpha power supporting auditory attention remains poorly understood. In the current electroencephalography (EEG) study, middle-aged and older adults (*N* = 29; ~40-70 years) performed a dichotic listening task that simulates a challenging, multi-talker scenario. We examined the extent to which the modulation of 8-12 Hz alpha power would serve as neural marker of listening success across age. With respect to the increase in inter-individual variability with age, we examined an extensive battery of behavioral, perceptual, and neural measures. Similar to findings on younger adults, middle-aged and older listeners′ auditory spatial attention induced robust lateralization of alpha power, which synchronized with the speech rate. Notably, the observed relationship between this alpha lateralization and task performance did not co-vary with age. Instead, task performance was strongly related to an individual’s attentional and working memory capacity. Multivariate analyses revealed a separation of neural and behavioral variables independent of age. Our results suggest that in age-varying samples as the present one, the lateralization of alpha power is neither a sufficient nor necessary neural strategy for an individual’s auditory spatial attention, as higher age might come with increased use of alternative, compensatory mechanisms. Our findings emphasize that explaining inter-individual variability will be key to understanding the role of alpha oscillations in auditory attention in the aging listener.

## Introduction

Selective attention allows us to separate and filter relevant from irrelevant information and thus supports behavior in many real-life situations. Everyday communication, for example, is often complicated by the temporal concurrence of different auditory signals. Here, selective auditory attention aids speech perception and comprehension by tuning out distracting sound sources like background noise or competing talkers (e.g., the ‘cocktail party problem’; Cherry, 1953; McDermott, 2009) to enable a specific focus on behaviorally relevant speech.

With respect to the neural implementation, there is a growing body of evidence that strengthens the functional link between neural oscillations and mechanisms of selective attention across different modalities (e.g., Foxe *et al.*, 1998; Klimesch, 1999; Fries *et al.*, 2001; Lakatos *et al.*, 2008; 2013). Within this line of research, modulations of oscillatory power in the 8-12 Hz alpha band have emerged as a candidate mechanism thought to underpin selective inhibition of neural activity involved in the processing of to-be-ignored signals (Jensen & Mazaheri, 2010; Strauß *et al.*, 2014). Importantly, numerous studies targeting different modalities and cognitive domains have shown that spatially directed attention leads to a corresponding hemispheric lateralization of oscillatory alpha-band activity with relative increases in power in the ipsilateral hemisphere and concurrent decreases in the contralateral hemisphere (visual: Worden *et al.*, 2000; Kelly *et al.*, 2006; Thut, 2006; Rihs *et al.*, 2007; Händel *et al.*, 2011; somatosensory: Haegens *et al.*, 2011; Bauer *et al.*, 2012; auditory: Kerlin *et al.*, 2010; Banerjee *et al.*, 2011; Müller & Weisz, 2011; Ahveninen *et al.*, 2013; Frey *et al.*, 2014).

However, most of the available evidence on the role of lateralized alpha power in the service of selective attention stems from research focused on the visual modality. Although visual and auditory attention appear to be under control of similar basic neural mechanisms (Shinn-Cunningham, 2008), findings from the visual domain do not necessarily represent direct analogues of the neural implementation of spatial attention in the auditory domain. The inherent temporal nature of auditory signals such as human speech requires attentional mechanisms that can flexibly adapt to its temporal dynamics. In fact, a recent study from our laboratory (Wöstmann *et al.*, 2016) has shown that the modulation of lateralized alpha power in response to a dichotic listening task aligns not only spatially, but also temporally to the attended speech input. Crucially, a more pronounced attentional modulation of alpha power correlated with increased listening success.

Aging listeners are known to experience difficulties in multi-talker listening situations, but it is unclear whether these difficulties are based on declining auditory perceptual acuity (Fostick *et al.*, 2013), decline of cognitive functioning, or both (Wingfield *et al.*, 2005; Anderson *et al.*, 2013). However, even less is known about age-related changes in the neural mechanisms of auditory spatial attention and how such changes may impact behavior. EEG studies focusing on event-related potentials (ERPs) have investigated age-related changes in auditory spatial attention using either dichotic listening or cued-attention tasks (Ford *et al.*, 1979; Woods, 1992; Karayanidis *et al.*, 1995; Gaeta *et al.*, 2003; Bennett *et al.*, 2004; Getzmann & Falkenstein, 2011; Getzmann, 2012; Getzmann *et al.*, 2016; for a recent review see Zanto & Gazzaley, 2014). The overall pattern of results emerging from these studies points towards an age-related delay in the onset of several early ERP components including the P2 and contingent negative variation (CNV), as well as changes in the amplitude of the P2, P3a, P3b and mismatch negativity. Studies that compared low- and high-performing individuals across age groups have found increases in amplitude in the P2-N2 complex or P3a component to be positively correlated with behavior in older but not younger adults (Getzmann & Falkenstein, 2011; Getzmann, 2012; Getzmann *et al.*, 2015). Taken together, these ERP results are generally in line with accounts suggesting that increasing age is accompanied by a slowing in processing speed and the recruitment of additional resources as a compensatory strategy (Cabeza *et al.*, 2002; Grady, 2012).

To date, all of the available evidence on changes in alpha lateralization with age comes from visual studies that contrasted young (~18-30 years) to older (~60-80 years) adults (Sander *et al.*, 2012; Hong *et al.*, 2015; Leenders *et al.*, 2016; but see Mok *et al.*, 2016) while virtually nothing is known about the period where one might suspect age-related change to first surface, that is, middle adulthood (~40-60 years) (Raz, 2005). In light of the well-attested fact that middle-aged adults already show signs of a gradual, if subtle, decline in cognitive abilities and sensory fidelity including hearing (Lin, 2011; Fabiani, 2012), and with new results emerging on the intricate interplay of hearing abilities and maintenance of cognitive fitness (Lin *et al.*, 2011; Wayne & Johnsrude, 2015; Deal *et al.*, 2016), targeting this age group should be a prime goal for the neuroscience of attention.

In the present study, we aim at closing this gap by investigating the neural dynamics of auditory spatial attention in an age group that specifically includes middle-aged adults but also encompasses older adults in the more typically studied age range. Thus, instead of relying on a comparison of younger and older age groups, we focus on uncovering continuous changes in behavioral and/or neural measures within our age-varying sample. We believe that the age range of 40-70 years presents an ideal sample to investigate how even subtle age-induced changes in sensory, cognitive, or neural processes may alter the ways in which they work together to support successful behavior.

We used an adapted version of the dichotic listening paradigm established by Wöstmann *et al.* (2016) that has been shown to reliably tap into the neural implementation of auditory spatial attention in time and space. In this task, participants were instructed to pay attention to one of two dichotically presented streams of spoken numbers and to then report the attended numbers as accurately as possible. While success on this task does not only rely on the implementation of spatial attention but e.g., also on short-term memory, it is still the most important prerequisite as performance critically hinges on the ongoing separation of speech streams by selective amplification and attenuation.

As highlighted above, for younger adults, Wöstmann *et al.* (2016) found a link between the attentional modulation of lateralized alpha power, thought to implement auditory spatial attention, and task performance. With respect to our sample of middle-aged and older adults, we were thus asking whether the fidelity of alpha power lateralization tuned to the rhythm of speech could serve as a neural marker of listening success in the aging adult. Importantly, in answering this main research question, we did not necessarily rely on finding a pronounced decrease in task performance with increased age. It is a well-attested phenomenon that older adults may achieve similar levels of task performance as younger adults but they do so by relying on different cognitive and neural strategies. It is precisely this change in the link between brain and behavior that is at the heart of the present study.

To investigate this matter, we did not only focus on the neural measures and behavioral results derived from this task but took a more comprehensive approach to characterize and understand the listening behavior in middle-aged and older adults. We followed this approach to account for the observation that the degree to which aging individuals show neural degradation or cognitive decline is accompanied by a considerable degree of inter-individual variability that increases with age (Rapp & Amaral, 1992; Li *et al.*, 2001; Hedden & Gabrieli, 2004; Fabiani, 2012). We thus included a variety of tests and questionnaires covering different domains, including attention, working memory, verbal intelligence and personality traits, as well as measures assessing quality of hearing in an objective and subjective manner.

With more specific evidence on selective auditory attention in middle-aged or older adults missing, our hypotheses were based on the evidence available from younger adults who performed the same task, and on results from studies investigating age-related effects on the lateralization of alpha power in the context of visual working memory. Collectively, these results suggest that increasing age correlates with a less pronounced or even absent lateralization of alpha power (Sander *et al.*, 2012; Hong *et al.*, 2015; Leenders *et al.*, 2016). Further evidence shows that younger and older participants also differ in their ability to uphold a heightened level of overall alpha power (Wöstmann *et al.*, 2015a; Henry *et al.*, 2017). We thus expected the strength and the sustenance of lateralized alpha power, and thereby possibly also its synchronization with speech, to decrease with age. Given the suggested link of attentional modulation of alpha power and behavior in younger adults, we expected this hypothesized change at the neural level to be reflected in poorer task performance as well. Finally, using a multivariate look at the full set of behavioral, perceptual, and neural measures, we aimed to uncover the multivariate patterns that characterize the link between behavioral and neural dynamics in the aging listener.

## Materials and methods

### Participants

Twenty-nine right-handed German native speakers (median age 56 years; range 39-69 years; 17 females; see Fig. 1A for age distribution) were included in the sample. Handedness was assessed using a translated version of the Edinburgh Handedness Inventory (Oldfield, 1971). All participants had normal or corrected-to-normal vision, did not report any neurological, psychiatric, or other disorders and were screened for mild cognitive impairment using the German version of the 6-Item Cognitive Impairment Test (6CIT; Jefferies & Gale, 2013). Data of four additional participants were excluded from all analyses; three due to technical problems during data acquisition and/or excessive artifacts and one due to overall unusually low (42 % hits; 44 % stream confusions) behavioral performance. Two participants dropped out before the EEG recording session and were thus excluded from all analyses as well. Participants gave written informed consent and received financial compensation. Procedures were approved by the ethics committee of the University of Lübeck and conformed to the Declaration of Helsinki.

**Figure 1.**
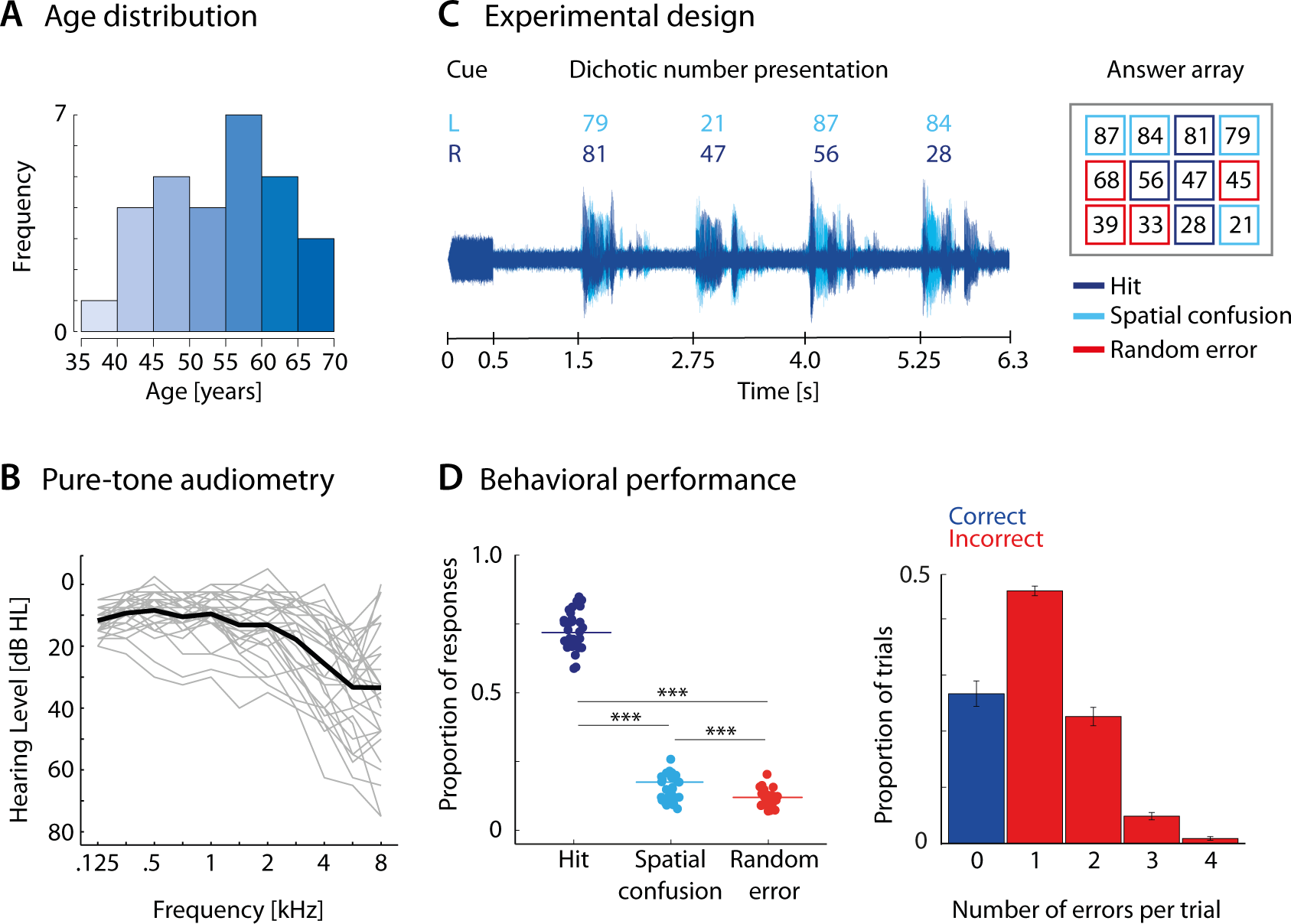
(A) Histogram showing age distribution of participants across seven 5-year age bins. (B) Air conduction hearing thresholds shown averaged across the left and right ear. Thick black line shows the average across all 29 participants; thin grey lines represent hearing curves of individual participants. Only participants with age-adequate hearing acuity were included. (C) Participants listened to two concurrent number streams. A cue tone played to one ear indicated the to-be-attended speech stream on a trial-by-trials basis. Participants selected the to-be-attended numbers from a visual answer array. Responses were categorized into hits (numbers selected from the to-be-attended stream), spatial confusions (numbers selected from the distractor stream) and random errors (numbers selected that were not presented in either of the streams). (D, left) Distribution of proportion of responses shown per response category. Dots show individual participants, colored horizontal lines represent the mean proportion per category across *N* = 29 participants. Hits occurred significantly more often than both spatial confusions and random errors. Spatial confusions in turn occurred more often than random errors. (D, right) Bar graphs display the average proportion of trials as a function of the number of errors per trial. Error bars indicate the standard error of the mean. Trials without any error were classified as correct (blue), trials with one or more errors as incorrect (red). [Color figure can be viewed at wileyonlinelibrary.com]

### Experimental sessions

Participants underwent two separate experimental sessions that were carried out on separate days for all but one participant (median difference between sessions: 4 days, range: 0-50 days). The first session consisted of a general screening procedure, detailed audiometric measurements, and a battery of cognitive tests and personality profiling. The order of tests and questionnaires administered in this session was kept constant for all participants. In the second session, we recorded the participants’ electroencephalogram (EEG) while they performed a dichotic listening task.

## Session 1 (Screening)

### Hearing acuity

Participants underwent audiometric testing that followed standard procedures (ISO 8253-1/3) and was carried out by a trained audiologist. Following otoscopic examination, pure tone hearing thresholds (air conduction: 0.125, 0.25, 0.5, 0.75, 1, 1.5, 2, 3, 4, 6, and 8 kHz; bone conduction: 0.25, 0.5, 1, 2, 3, 4, and 6 kHz) as well as uncomfortable loudness levels (0.5, 1, 2, 4 kHz) were measured separately for each ear in steps of 5 dB hearing level (HL). In addition, speech audiometry was assesses using the Freiburger speech intelligibility test with monosyllabic words presented at 65 dB sound pressure level (SPL) in quiet. Only participants with normal hearing or age-adequate mild-to-moderate hearing loss were included. Our definition of mild-to-moderate hearing loss was largely in agreement with the grades of hearing impairment defined by the World Health Organization (WHO) (mild: 26-40 dB HL, moderate: 41-60 dB HL). However, we did not exclude participants who had hearing threshold elevated to >60 dB HL at higher frequencies (4 kHz or above), as long as they had hearing thresholds <30 dB at 1 kHz and speech intelligibility scores of >80 % at 65 dB SPL. Additional exclusion criteria included interaural asymmetries of more than 20 dB in the pure-tone average (PTA, including the primary speech frequencies of 0.5, 1, 2, and 4 kHz), and hearing loss profiles not following the typical age-related gradual decline at higher frequencies (e.g., increased thresholds restricted to lower or mid-range frequencies, or observed uniformly across all frequencies. Air conduction hearing thresholds of all participants along with the group average are presented in Figure 1B.

To obtain a measure of self-perceived hearing (dis-)abilities, we administered a shortened version of the Spatial, Speech and Hearing Quality Scale (SSQ; Gatehouse & Noble, 2009). This assessment tool describes a number of real-life listening scenarios with a particular emphasis on speech and spatial hearing including the separation of an attended talker from either competing talkers or background noise. Participants rated their relative hearing difficulty in these situations on a scale from 0 to 10 with smaller numbers indicating more difficulty.

### Attention

Following audiometric testing, participants performed a standard paper-and-pencil test measuring their attentional capacity (d2-R; Brickenkamp *et al.*, 2010). In this test, participants were asked to cancel visual target letters in a list of highly similar distractor letters. We utilized two separate, standardized measures from this test: processing speed (i.e., sum of processed target items) and concentration performance (i.e., the sum of correctly marked target items minus the sum of incorrectly cancelled distractors; Bates & Lemay, 2004).

### Working memory

Working memory was measured using the auditory forward and backward digit span test (part of the WAIS-IV; Wechsler, 2008). Participants listened to lists of digits between 1 and 9 that increased in length. After presentation participants were asked to immediately repeat the digits either in their order of presentation (forward digit span) or in reverse order starting with the last digit (backward digit span). Forward digit span was assessed first.

### Verbal intelligence

To obtain a marker of crystallized intelligence, we carried out the German multiple-choice vocabulary intelligence test (MWT-B) that has been shown to strongly correlate with verbal and global intelligence scores of other standard tests and often increases with age (Lehrl *et al.*, 1995). Participants had to perform lexical decisions on 37 items, i.e., they had to spot an existing word among non-word distractors.

### Personality traits

Finally, we collected information on the participants’ core personality dimensions using the German 10-item short version of the Big Five Inventory (BFI-10; Rammstedt & John, 2007). Participants rated themselves on two items for each of the Big Five personality dimensions (extraversion, agreeableness, openness, neuroticism and conscientiousness).

The first experimental session lasted on average about 1.5 hours. A summary of the average performance on these tests together with basic demographic information is presented in Table 1.

**Table 1.**
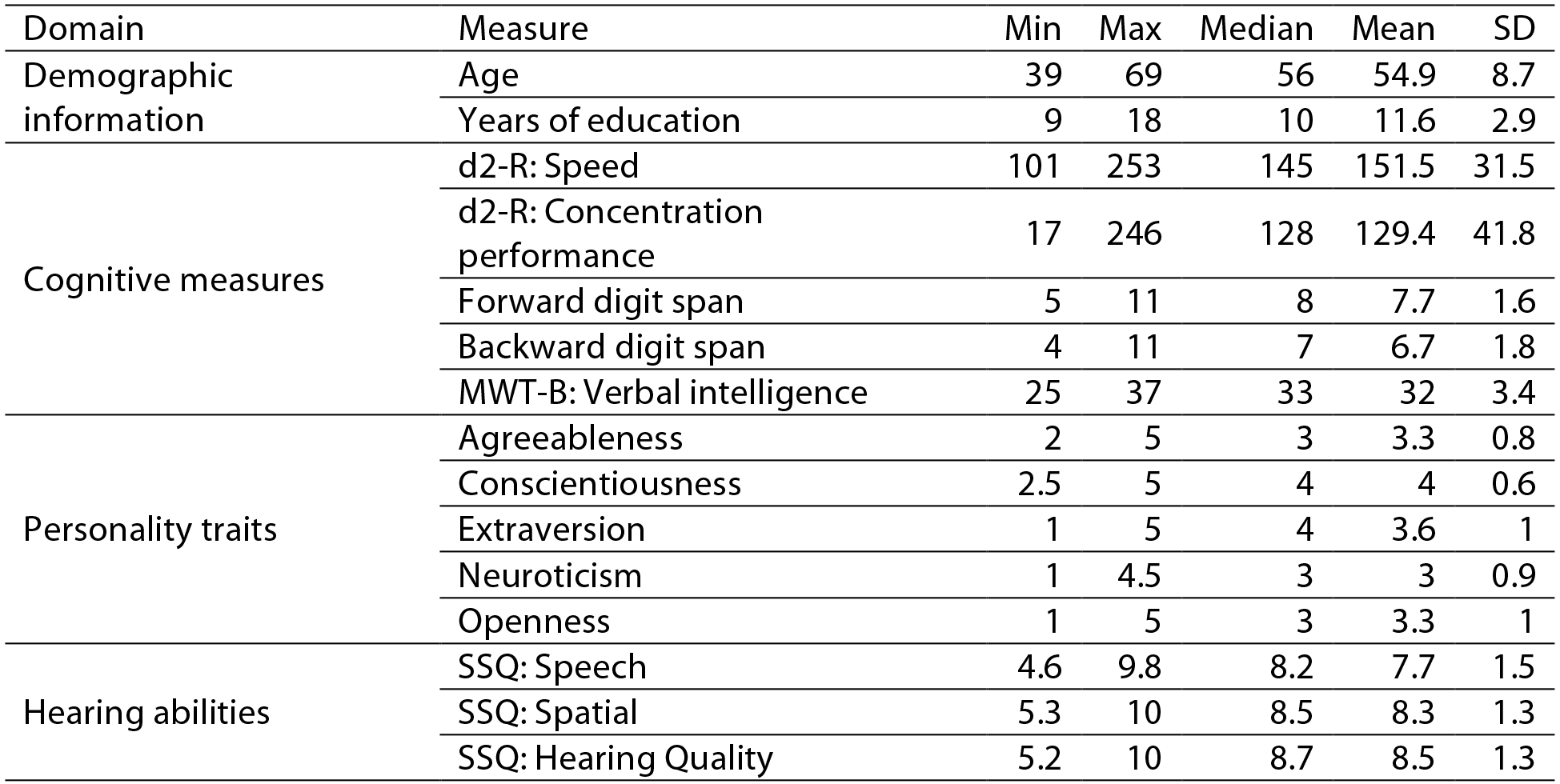
Descriptive statistics of behavioral and demographic variables.

## Session 2 (EEG experiment)

### Stimuli

Auditory stimuli consisted of a 1000-Hz pure tone (500 ms duration, 50 ms linear onset ramp) and the recordings of German double-digit numbers that were adopted from previous studies (Wöstmann *et al.*, 2015a; Wöstmann *et al.*, 2015b; Wöstmann *et al.*, 2016). All four-syllable numbers between 21 and 99 (i.e., excluding integer multiples of ten) were recorded at a sampling rate of 44.1 kHz and digitally adapted to have an average duration of ~1 s. Root mean square (rms) intensity (−22 dB Full Scale, FS) was equalized across numbers and the 1000-Hz pure tone.

### Experimental design and procedure

The dichotic listening task used in this study is an adapted version of the paradigm by Wöstmann *et al.* (2016; see for full details). In this task, participants listened to two speech streams consisting of four double-digit numbers that were presented concurrently to the left and right ear. The perceptual centers of each pair of simultaneously presented numbers were temporally aligned to achieve a tightly synchronized onset across the left and right ear. The first and second digit of two simultaneously presented numbers were always distinct. Spatial cue and numbers were masked by continuous white noise at a signal-to-noise ratio of+10 dB.

A schematic representation of the experimental paradigm is shown in Figure 1C. Each trial started with the presentation of a fixation cross in the middle of a 24” touch screen (ViewSonic TD2420) positioned within an arm′s length. A 500-ms spatial cue tone played to either the left or right ear signaled the to-be-attended number stream on a trial-by-trial basis. After a subsequent anticipation period of 1000 ms, the two spoken number streams were presented. The onsets of two consecutive numbers were 1.25 s apart, resulting in a presentation rate of 0.8 Hz. Note that the slightly faster presentation rate was the only change in the task design compared to the version used by Wöstmann *et al.* (2016) who presented the dichotic numbers at a speech rate of 0.67 Hz. Following a short retention period of approximately one second (jittered between 0.8-1.2 s), a response array was shown in the middle of the screen. Participants were instructed to select the four numbers presented on the to-be-attended side in any order using the touch screen. The reason for using a touch screen instead of a computer mouse was to reduce age-related influences in handling the response device on performance. Among the twelve displayed numbers were the four target numbers from the to-be-attended side, four distractor numbers from the to-be-ignored side and four random numbers not presented in any stream. Once the last of four numbers had been chosen on the screen, the next trial started after a short jittered inter-trial interval (mean: 2.5 s; range: 2.2-2.8 s).

Following instructions, participants performed a few practice trials to familiarize themselves with the dichotic listening task. To account for differences in hearing acuity within our group of participants, individual hearing thresholds for a 500-ms fragment of the dichotic number stimuli were measured using the method of limits. All stimuli were presented 50 dB above the individual sensation level using Sennheiser HD-25 headphones. During the experiment, each participant performed five blocks of 30 trials for a total of 150 trials. Participants took short breaks between blocks. The order of trials was fully randomized with the constraint that each block presented equal numbers of attend-left and attend-right trials. Including preparation time, the second session lasted approximately 2.5 to 3 hours.

### Behavioral data analyses

We evaluated participants′ behavioral performance in the dichotic listening task by grouping responses into three categories: correctly selected numbers from the to-be-attended stream were classified as *hits*, selected numbers that instead appeared on the to-be-ignored stream as *spatial confusions*, and selected numbers that did not appear during a given trial as *random errors*. Accuracy was assessed by calculating the proportion of hits, spatial confusions and random errors per participant. For statistical analysis, we first ensured that the assumption of normality was not violated for any of the three response distributions (Shapiro-Wilk test; all *Ps* > 0.05). We then proceeded by conducting a one-way repeated-measures ANOVA with response category as within-subject factor and performed post hoc paired *t*-tests. For repeated-measures ANOVAs with more than one degree of freedom in the numerator, we report Greenhouse-Geisser (GG) epsilon and the GG-corrected *p*-values when sphericity was violated (Mauchly’s test). All behavioral analyses were performed using R (R Core Team, 2017).

### EEG recording and preprocessing

Participants were seated comfortably in a dimly-lit, sound-attenuated recording booth where we recorded their EEG from 64 active electrodes mounted to an elastic cap (Ag/AgCl; ActiCap / ActiChamp, Brain Products, Gilching, Germany). As part of the EEG preparation, we measured the distance between the left and right preauricular points, as well as between the nasion and inion and placed the vertex electrode CZ at their intersection. Electrode impedances were kept below 5 kΩ. The signal was recorded with a passband of DC to 280 Hz, digitized at a sampling rate of 1000 Hz and referenced on-line to the left mastoid electrode (TP9, ground: AFz).

For subsequent off-line EEG data analyses, we used the Fieldtrip toolbox (version 2016-06-13; Oostenveld *et al.*, 2011) for Matlab (version R2016a, The MathWorks, Inc., Natick, US) and customized Matlab scripts. The continuous EEG data were highpass-filtered at 0.3 Hz (finite impulse response (FIR) filter, zero-phase lag, order 5574, hanning window) and lowpass-filtered at 180 Hz (FIR filter, zero-phase lag, order 100, hamming window). We segmented the continuous data into epochs of −2 to 10 s relative to the onset of the trial-initial cue tone to capture the entire auditory stimulation interval. Independent component analysis (ICA) using Fieldtrip′s default runica algorithm was used to remove eye blinks and lateral eye movements, muscle activity, heartbeats and channel noise. The classification into brain and artefact components was based on their topographical, as well as spectral representations. Since we were only interested in frequencies up to 20 Hz, this procedure included the removal all components that were dominated by high-frequency (i.e. > 30 Hz) noise. Following ICA, we ran an automated rejection algorithm that removed trials in which the amplitude of any individual EEG channel exceeded a range of 200 μV. Finally, EEG data were re-referenced to the average of all EEG channels (average reference). On average, 57.6 % (± 9 %) of components and 3.8 % (± 3 %) of trials were removed. The number of removed components and trials did not correlate with age (*Ps* > 0.05).

## Key neural measures of interest

The analysis of EEG data was constrained *a priori* by following as closely as possible the analysis procedure carried out by Wöstmann *et al.* (2016). We focused on the following key neural measures of interest. We analyzed inter-trial phase coherence (ITPC) in slow cortical oscillations (1-5 Hz) as a reflection of more bottom-up driven sensory processing of speech. The investigation of top-down driven attentional modulation of neural dynamics in space and time focused on the strength and modulation of the lateralization of 8-12 Hz alpha power across the trial. Analyses on the extent to which lateralized alpha power (or low-frequency ITPC) would synchronize with the presented speech, and whether the strength of this synchronization would be predictive of behavior examined the modulation of these neural measures at the speech rate of 0.8 Hz. Since Wöstmann and colleagues reported a lateralization of alpha power that was not only temporally but also spatially specific, i.e., encompassing not only supramodal but also auditory regions, we additionally investigated the involvement of different brain regions via reconstruction of the neural sources. The details of the individual analysis steps are described below.

### Time-frequency analyses of oscillatory dynamics

For time-frequency representations of single-trial EEG oscillatory power and phase, we convolved individual epochs ranging from −1.5 to 9 s with a family of Morlet wavelets and obtained complex wavelet coefficients for frequencies of 1-20 Hz with a frequency precision of 0.5 Hz and a temporal precision of 0.05 s. To analyze inter-trial phase coherence (ITPC), we used a wavelet width of 3 cycles while total power calculations were performed on Fourier coefficients obtained with a wavelet width of 7 cycles. ITPC was defined as the magnitude of amplitude-normalized complex wavelet coefficients that were averaged across single trials. Oscillatory power representations were calculated by squaring the magnitude of the estimated complex-valued Fourier coefficients.

To visualize dynamics in oscillatory power across all frequencies during the presentation of cue tone and numbers (0-6.5 s), we calculated power changes by subtracting and dividing power averaged across trials by the mean power in this interval (relative change baseline). Note that we did not use a pre-stimulus time window as baseline because anticipatory effects led to a disproportionally strong increase in absolute 8-12 Hz alpha power before the presentation of the cue tone. Grand average representations of ITPC and oscillatory power were obtained by averaging across all 29 participants.

### Attentional modulation indices

In line with the analyses reported in Wöstmann *et al.* (2016), we calculated two indices that describe attentional modulation of neural responses: the attentional modulation index (AMI) and the alpha lateralization index (ALI; Haegens *et al.*, 2011). The AMI, calculated per channel for both 1-5 Hz ITPC (AMI_ITPC_) and 8-12 Hz absolute alpha power (AMI_α_), offers a spatial representation of attentional effects on these two neural measures and is derived as follows: AMI = (attend left - attend right) / (attend left + attend right). Positive AMI values for a dependent measure (ITPC or alpha power, respectively) reflect higher levels during attend-left compared to attend-right trials. The reverse relation holds for negative values. We analyzed the topological distribution of these values separately for ITPC and alpha power in three time windows of interest (cue: 0-0.5 s, anticipation: 0.5-1.5 s, and number presentation: 1.5-6.3 s). Lateralization in each time window was statistically assessed by averaging AMI_α_ or AMI_ITPC_ values across all 15 posterior channels per hemisphere (excluding midline electrodes) and submitting the results to paired t-tests or Wilcoxon signed-rank tests when normality assumptions were violated. For intervals with a significant hemispheric lateralization at the group level, we also quantified the strength of the lateralization at the level of the individual participant and submitted these to the multivariate analysis of neural and behavioral measures (see below). To this end, we calculated the difference between mean AMI values across the same 15 left and right posterior channels where larger difference scores indicated a stronger lateralization.

We then selected for each participant the ten channels with the largest positive AMI_α_ values over the left posterior hemisphere and the ten channels with the largest negative AMI_α_ values over the right posterior hemisphere. For each of the two attention conditions (attend left versus attend right), the selected channels were classified as being either ipsi- or contralateral to the focus of attention. To obtain a time-resolved measure of attentional modulation of alpha power, we calculated the ALI [ALI_α_= (α_att_ipsi_-α_att_contra_)/(α_att_ipsi_+α_att_contra_)] over the time course of the entire trial using a 250-ms sliding window (Mesgarani & Chang, 2012).

To examine the degree to which fluctuations in 1-5 Hz ITPC or 8-12 Hz ALI synchronized with the presented speech, we first averaged ITPC values across a cluster of eleven fronto-central electrodes (F1, F2, FC3, FC1, FC2, FC3, FC4, C1, C2, C3, C4) where phase locking in response to auditory input tends to be most pronounced. We then calculated the Fast Fourier Transforms for the time courses of ITPC and ALI during number presentation (1.5-6.3 s; using zero-padding to obtain a frequency resolution of 0.01 Hz). The magnitude of the resulting complex coefficients was used as measure of frequency-specific modulation strength. In addition, to investigate the temporal relationship of ITPC and lateralized alpha power, we extracted the phase of each measure at the frequency of 0.8 Hz and quantified the concentration of phase angles across participants, as well as the average delay in phase between ITPC and ALI. To examine the impact of speech-synchronized modulation of lateralized alpha power on behavior, we contrasted the modulation depth of ALI at 0.8 Hz for correct and incorrect trials across individual participants, and tested for changes with age using mixed-effects modelling.

### EEG source localization

Source localization was carried out by applying the Dynamic Imaging of Coherent Sources (DICS) beamformer approach (Gross *et al.*, 2001) implemented in the Fieldtrip toolbox. To this end, we used a standard headmodel (Boundary Element Method, BEM; 3-shell) to calculate a common leadfield for a grid of 1 cm resolution. We then estimated the neural sources for three different measures of interest: auditory activation as reflected by 1-5-Hz ITPC, and the significant lateralization of alpha power and ITPC (reflected by AMI_α_ and AMI_ITPC_, respectively) in different intervals of the trial.

For the localization of auditory activation during cue and number presentation (time window: 0-6.3 s relative to cue onset), we calculated an adaptive spatial filter from the leadfield and the cross-spectral density of Fourier Transforms centered at 3 Hz with a ± 2 Hz spectral smoothing. This filter was applied to single-trial Fourier Transforms (1-5 Hz, in steps of ~0.15 Hz). ITPC at each grid point and frequency was calculated and averaged across frequencies. In addition, to localize the lateralization of ITPC during the cue interval of 0-0.5 s, we first estimated a common spatial filter restricted to this time window based on all trials. Next, we calculated the cross-spectral density of Fourier Transforms using the same parameters as above but separately for attend-left and attend-right trials. The common filter was then separately applied for source projection of the attend-left and attend-right conditions. At the source level, AMI_ITPC_ was calculated for each participant at each grid point.

The source localization of the alpha power attentional modulation index proceeded in a similar manner. Here, we estimated the cross-spectral density of Fourier Transforms centered at 10 Hz with ±2 Hz spectral smoothing separately for each attention condition and in all three time windows of interest (cue: 0-0.5 s, anticipation: 0.5-1.5 s, and number presentation: 1.5-6.3 s). Again, we built a common filter for each time window using the leadfield and cross-spectral density based on all trials that was then applied separately for the source projection of the attend-left and the attend-right condition. Lastly, AMI_α_ in source space was calculated for each participant at each grid point.

### Power spectral density

For our multivariate analyses (see below) we further included an estimate of the individual 1/f noise in the EEG, a neural measure that is thought to capture trait-like changes in the neural variability and the balance of neural excitation and inhibition across age (Voytek *et al.*, 2015; Gao *et al.*, 2017; Waschke *et al.*, 2017). As such it represents a task-independent measure of age-related changes in neural dynamics. We calculated the power spectral density (PSD) using participants′ single trials with a 2-s window slid across the trial with 50 % overlap. PSD estimates were then averaged across trials. For frequencies between 2-45 Hz, we estimated the linear fit to the power spectrum (excluding the 8-12 Hz alpha range; Voytek *et al.*, 2015) for each participant, and included the slope of the linear regression line at electrode POz as a variable in the multivariate analyses.

## Multivariate analyses of neural and behavioral measures

To explore and visualize the complex relationship between performance in the dichotic listening task, age, years of education, as well as the set of behavioral and neural measures, we calculated the correlation matrix between these variables. We used Spearman rank correlation for pairs of linear measures and circular-linear correlations for pairs of circular and linear variables, respectively. We included the average scores of the shortened Big Five inventory as personality traits, and speed, concentration performance, verbal intelligence, and forward and backward digits span as cognitive measures. To describe the fidelity of auditory perception, we included as an objective measure of hearing acuity the pure-tone average across both ears, and as a subjective measure of hearing abilities the average scores for each of the three subdomains of the SSQ questionnaire. Lastly, as neural measures we included the strength of the AMI_α /ITPC_ for intervals showing significant hemispheric lateralization, the amplitude of the ALI and ITPC at 0.8 Hz, the phase delay at 0.8 Hz for the two measures, as well as the slope of the power spectral density.

We corrected for multiple comparisons by controlling the false discovery rate (FDR; Benjamini & Hochberg, 1995) that was set to 10 % in this exploratory analysis. Finally, we performed principal component analysis (PCA) on the same set of variables (scaled to unit variance) to further investigate the link between (age-related) changes in neural dynamics, task-performance and cognitive abilities by exploring the internal structure of this complex data set.

## Statistical testing and effect sizes

We applied parametric tests when the data conformed to normality assumptions (*p* > 0.05 in Shapiro-Wilk test) and appropriate non-parametric alternatives, including rank transformation, otherwise. For effect sizes, we report r_equivalent_ (bound between 0 and 1; Rosenthal, 1994; Rosenthal & Rubin, 2003) for *t*-tests and their non-parametric alternatives, as well as partial eta-squared for repeated-measures ANOVAs. For circular statistics, we report the circular-linear correlation of phase angles and condition labels. For circular Rayleigh tests, we report the resultant vector length (*r*).

For linear mixed-effects models, we followed an iterative model fitting procedure, which started with an intercept-only null model. Fixed effects terms were added in a step-wise procedure and the change of model fit (performed using maximum likelihood estimation) was assessed using likelihood ratio tests. Deviation coding was used for categorical predictors. We report *p*-values for individual model terms that were derived using the Satterthwaite approximation for degrees of freedom (Luke, 2017). Post hoc linear contrasts for predictors with more than two levels were carried out on predicted marginal means (‘least-square means’) and standard errors estimated from the model (Lenth, 2016). In the absence of a standard effect size measure for individual terms in linear mixed models, we report the mean difference between predictor levels.

To enhance the interpretability of non-significant effects including age in the present study, we also calculated the Bayes Factor (BF; using the *BayesFactor* package in R). In detail, when comparing two statistical models, the BF indicates how many times more likely the observed data are under the alternative, more complex compared to the simpler model. While the assignment of categorical boundaries is unnecessary as the Bayes Factor is directly interpretable as an odds ratio there is still considerable agreement that a BF <0.1 is interpreted as providing strong evidence in favor of the null hypothesis and a BF >10 as strong evidence against it (Rouder *et al.*, 2009). Note that this approach also helps to overcome some of the limitations associated with the comparably small sample size available here: using a Bayes-Factor-based testing approach is recommended to specifically circumvent questions of whether the sample was simply too small to detect an effect (Cousijn *et al.*, 2014; Wöstmann & Obleser, 2016).

## Results

### Middle-aged and older adults maintain high levels of attentional control

Figure 1D shows the overall behavioral performance on the dichotic listening task broken down by the three response categories. The average proportions of hits (mean ± SD 0.74 ± 0.08), spatial confusions (0.15 ± 0.05), and random errors (0.12 ± 0.03) differed significantly (*F*_*2,56*_ = 781.9; *ɛ* = 0.55; *p* < 0.001; *η*^2^_*p*_ = 0.97). Pair-wise comparisons revealed that the proportion of hits was significantly higher than the rate of spatial confusions (*t*_*28*_ = 25.8; *p* < 0.001; *r* = 0.98) and random errors (*t*_*28*_ = 32.1; *p* < 0.001; *r* = 0.99). Replicating the results of Wöstmann *et al.* (2016), spatial confusions occurred significantly more often than random errors (*t*_*28*_ = 4.7; *p* < 0.001; *r* = 0.66), highlighting the distracting effect of the to-be-ignored speech stream.

### Preserved attentional modulation of alpha power in middle-aged and older adults

The analysis of neural oscillatory dynamics focused on two key measures, low-frequency phase coherence and alpha power, and on how they varied in line with the attentional constraints imposed by the dichotic listening task.

Figure 2A presents changes in inter-trial phase coherence (ITPC) across all trials. As can be seen, the onsets of cue tone and individual numbers are temporally aligned with pronounced bursts of increased phase coherence in frequencies below 10 Hz. Neural sources of 1-5 Hz ITPC during auditory stimulation (0-6.3 s) were localized to superior and middle temporal (i.e., auditory) regions (Fig. 2B). Figure 2D shows changes in overall neural oscillatory power throughout the duration of the trial. Oscillatory power in the alpha and lower-beta band starts off strongest around the trial-initial cue tone, and then decreases to a level below baseline in a fluctuating manner.

**Figure 2.**
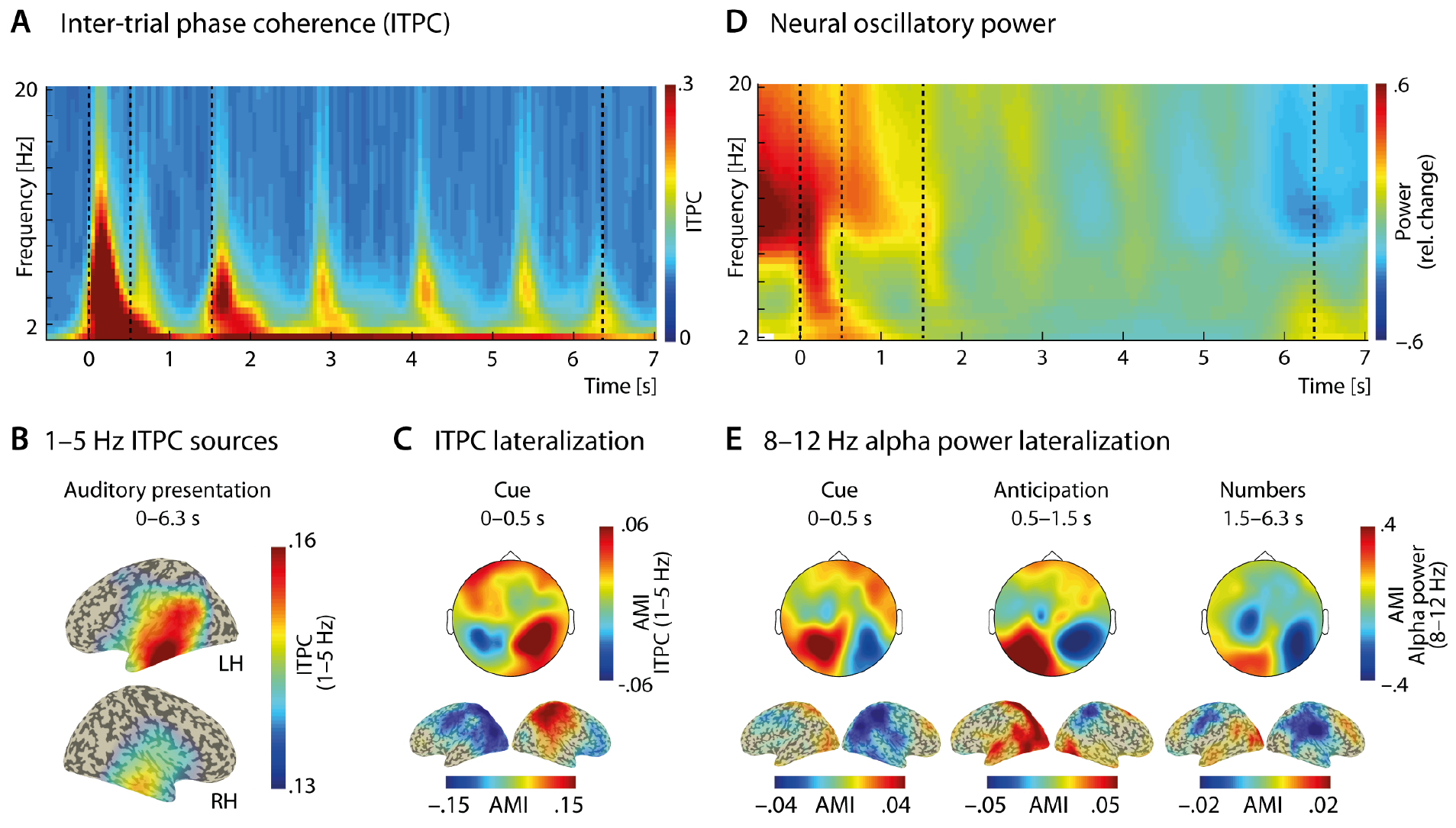
(A) Inter-trial phase coherence (ITPC) and (D) oscillatory power averaged across all trials, all 62 scalp electrodes (i.e., excluding mastoid electrodes) and *N* = 29 participants. Dotted vertical lines indicate the onset and offset of the spatial cue tone (0-0.5 s), anticipation interval (0.5-1.5 s), and number presentation (1.5-6.3 s). Power changes are calculated relative to a whole trial (0-6.5 s) baseline interval. (B) Neural sources of 1-5 Hz ITPC during cue tone and number presentation (0–6.3 s) were localized to auditory cortex region in the temporal lobes. ITPC values were projected onto an inflated brain model and thresholded to only show the highest 25 % of values. (C) Topographic maps and reconstructed neural sources shown for the attentional modulation index (AMI) for 1-5 Hz ITPC during the cue interval (0-0.5 s) and in (E) for 8-12 Hz alpha power during the cue (0-0.5 s), anticipation (0.5-1.5 s), and number presentation (1.5-6.3 s) interval. Warm colors indicate relatively stronger neural signals (ITPC or power) in attend-left compared to attend-right trials (and vice versa for cold colors). AMI values at the source level mirror the hemispheric lateralization at the sensor level and involve mostly occipital, superior and inferior parietal but also auditory regions in the temporal cortices. [Color figure can be viewed at wileyonlinelibrary.com]

We examined attention-related changes in ITPC and power, specifically in the 8-12 Hz alpha band, by means of the Attentional Modulation Index (AMI) that contrasts neural responses during attend-left and attend-right trials. As Figure 2C shows, we found a significant cue-evoked lateralization of AMI_ITPC_ only during the cue interval (0-0.5 s; *t*_*28*_ = 6.35; *p* < 0.001; *r* = 0.77), with larger mean AMI_ITPC_ values over the right compared to the left posterior hemisphere (see Fig. S1 for anticipation and number presentation intervals). The same pattern was found at the source level with the strongest positive AMI_ITPC_ values over the right superior and inferior parietal cortex and the strongest negative values over the left parietal and occipital cortex.

As expected, the distribution of AMI_α_ values described the reverse pattern with positive values over left posterior electrodes and negative values over right posterior electrodes (Fig. 2E). Here, the spatial attention-induced lateralization of 8-12 Hz alpha power was most pronounced during the anticipation interval (0.5-1.5 s; Wilcoxon signed-rank test; *z* = 3.81; *p* < 0.001; *r* = 0.5) but also present during the cue (0-0.5 s; Wilcoxon signed-rank test; *z* = 3.58; *p* < 0.001; *r* = 0.47) and number presentation (1.5-6.3 s; *t*_*28*_ = 3.25; *p* = 0.03; *r* = 0.52). Again, the spatial distributions of AMI_α_ at the scalp level were mirrored by comparable hemispheric differences at the source level. The largest positive and negative AMI_α_ values were clustered across the superior and inferior parietal and occipital cortex but also extending all the way to auditory regions in the superior temporal lobes.

### Temporal dynamics of lateralized alpha power and low-frequency phase coherence

Figure 3A presents the temporal relationship of lateralized alpha power and low-frequency phase coherence in reference to the rhythmic speech input. As first highlighted by Wöstmann and colleagues (2016), the dynamics of both lateralized alpha power, expressed by the Alpha Lateralization Index (ALI), and 1-5 Hz phase coherence were in their unique ways temporally coupled to the presentation rate of speech input (here: 0.8 Hz). While peaks in low-frequency ITPC were tightly aligned to the onsets of auditory stimulation, they were followed by peaks in lateralized alpha power that gradually decreased in amplitude over time. The difference in phase angles at the presentation rate of 0.8 Hz for ITPC versus ALI was statistically significant (Parametric Hotelling paired-sample test; mean delay: ~87°, or 302 ms; *F*_*54*_ = 18.80; *p* < 0.001; *r* = 0.67). At the same time, the concentration of phase angles across participants was significantly clustered for ITPC (Circular Rayleigh test; *z* = 23.80; *p* < 0.001; *r* = 0.91) but weaker and only approaching statistical significance for the ALI (Circular Rayleigh test; *z* = 2.68; *p* = 0.068; *r* = 0.30).

**Figure 3.**
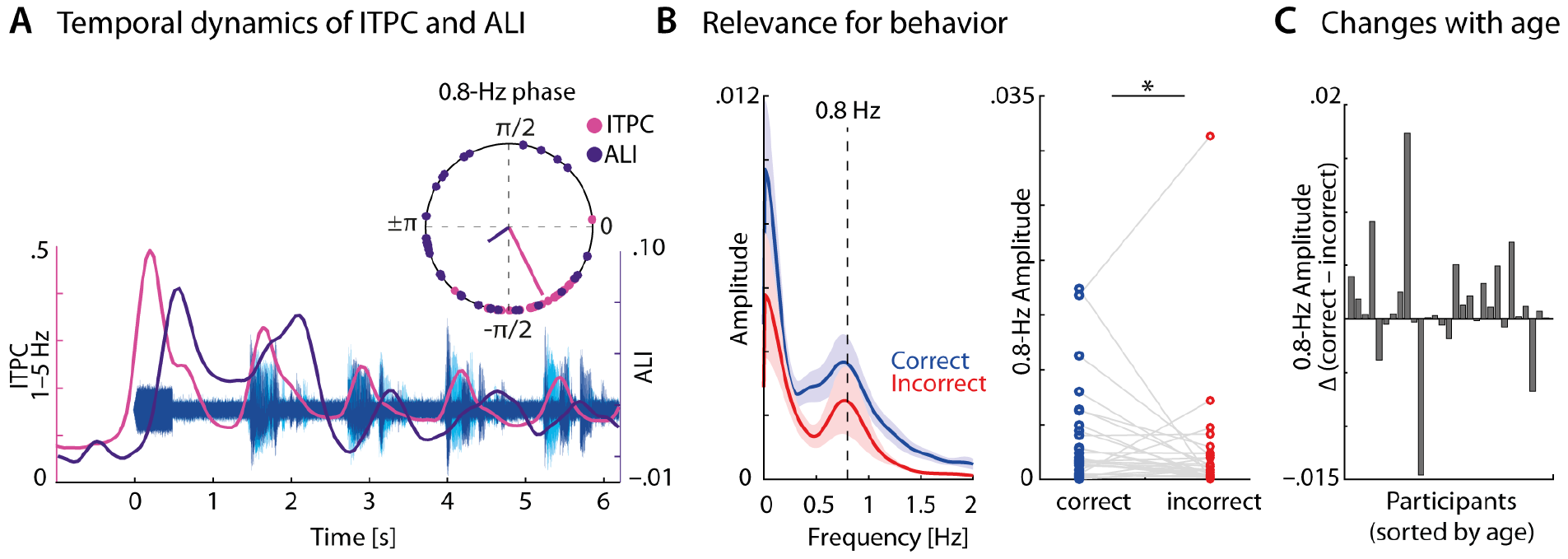
(A) 1-5 Hz inter-trial phase coherence (ITPC) averaged across fronto-central electrodes (F1, F2, FC3, FC1, FC2, FC3, FC4, C1, C2, C3, C4; pink) and average 8-12 Hz alpha lateralization index (ALI; purple) shown in temporal relation to auditory stimulus throughout the trial. Dichotic numbers were presented at a rate of 0.8 Hz. Inset shows individual phase angles of 0.8 Hz amplitude modulation of ITPC and ALI during number presentation (1.5-6.3 s). Colored lines show the mean phase angles across all 29 participants. (B, left) Amplitude spectra of ALI during number presentation shown separately for correct (zero errors; blue) and incorrect (one or more errors; red) trials. Dashed line indicates the number presentation rate of 0.8 Hz. Shaded error bands indicate ± 1 SEM. (B, right) The 0.8-Hz amplitude of ALI was significantly higher in correct compared to incorrect trials. * *p* < 0.05. (C) Bar graphs show the difference in 0.8-Hz amplitude of correct and incorrect trials for all 29 participants sorted by age (increasing from left to right). The difference in 0.8-Hz amplitude did not vary as a function of age. [Color figure can be viewed at wileyonlinelibrary.com]

### Age does not affect the relationship of alpha modulation and behavioral performance

We examined the relationship of the strength of 0.8-Hz alpha-lateralization (ALI) modulation and behavioral performance across age. To this end, we categorized trials in line with the approach taken by Wöstmann *et al.* (2016) who classified trials without any errors as correct and all other trials as incorrect (Fig 1C). Figure 3B shows amplitude spectra of the ALI estimated for the interval of number presentation (1.5-6.3 s) separately for correct and incorrect trials.

Most participants showed stronger 0.8-Hz ALI modulation for correct compared to incorrect trials (Fig. 3C). To test this difference in modulation statistically for all participants but also across age, we submitted rank-transformed amplitude values to a linear mixed-effects model fitting procedure. Model comparison revealed that the best-fitting linear mixed effects model included accuracy as categorical fixed factor (correct vs. incorrect; *t*_*28*_ = 2.12; *p* = 0.04; mean amplitude difference = 0.001) and participants as random factor. Notably, the inclusion of age or the interaction of accuracy x age did not improve model fit compared to the simpler model (*χ*^*2*^_*1*_ = 0.07; *p* = 0.8; *BF* = 0.32, and *χ*^*2*^_*1*_ = 0.83; *p* = 0.67; *BF* = 0.18, respectively).

### Inter-individual differences outweigh aging effects on auditory attention

Figure 4A shows the pattern of relationships that emerged from pair-wise correlations of our variables of interest. We found performance (i.e., hit rate) in the dichotic listening task to be best predicted by measures reflecting attentional capacity, namely speed and concentration performance, as well as by forward (and to a lesser degree backward) digit span. Notably, increasing age was only a weak predictor of task performance or of any of the derived neural measures, including the power spectral density (PSD) slope, but correlated more strongly with (increasing) verbal intelligence and (decreasing) hearing acuity. Figure 4B provides a detailed overview of the variables that were most strongly linked to age or performance on the task.

**Figure 4.**
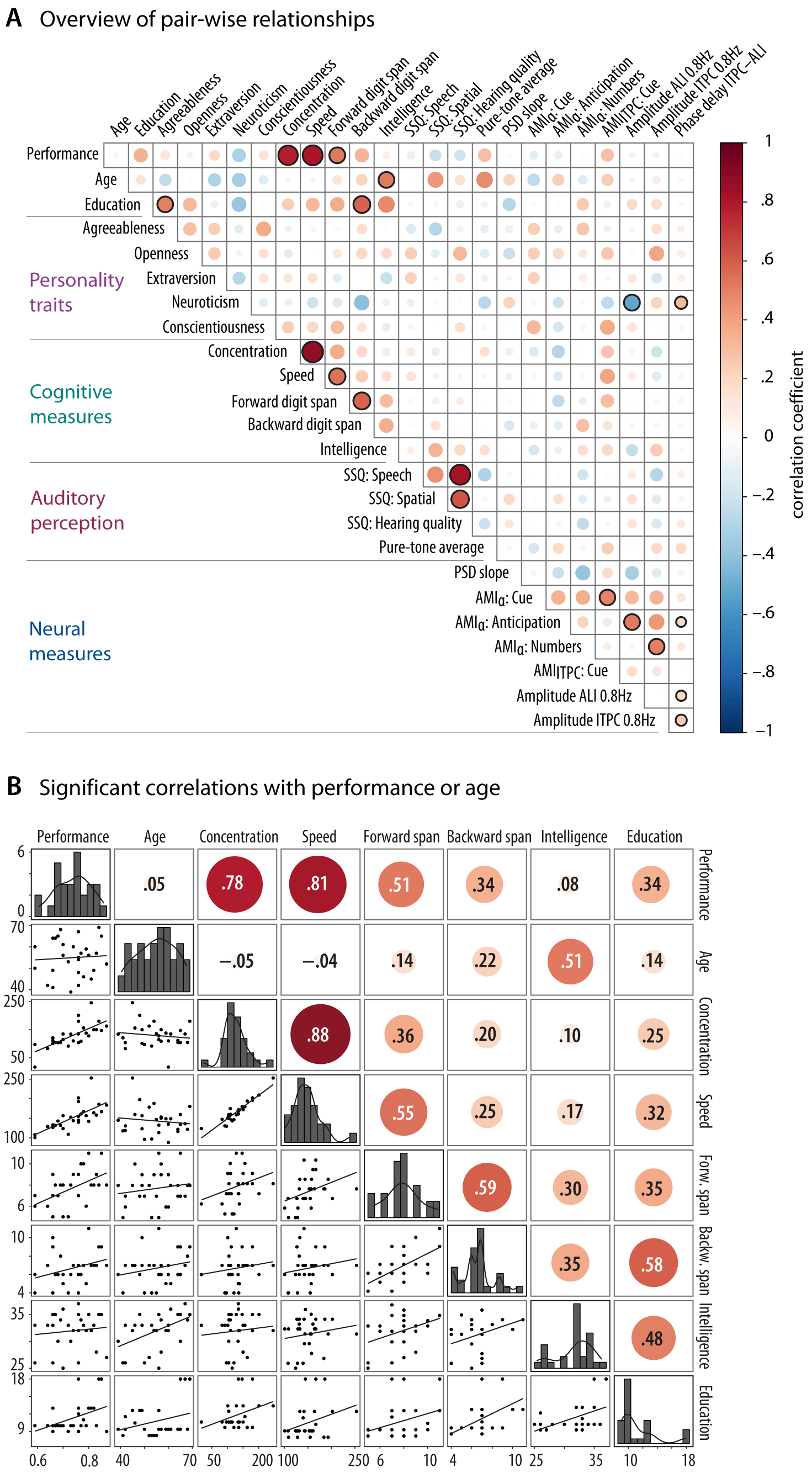
(A) Correlogram showing all pair-wise correlations between variables of interest (Spearman correlations for pairs of linear variables and circular-linear correlation for correlations involving the 0.8-Hz phase delay of ITPC and ALI). Color and size of circles indicate the strength of the relationship between two variables. Circles with black outline denote correlation coefficients significant after applied correction for multiple comparisons (FDR, q=0.1). Note that FDR correction was performed jointly for p-values resulting from Spearman and circular-linear correlations. Variables are spatially arranged to form five groups describing different domains. (B) Upper triangle: Pair-wise Spearman correlations of behavioral measures significantly correlated with either performance or age. Diagonal: Histograms along with density curve shown per variable. Lower triangle: Scatterplots with linear regression line. [Colour figure can be viewed at wileyonlinelibrary.com]

Figure 5A shows the projection of the examined variables into the vector space spanned by the first two principal components derived from principal component analysis (PCA). Together, these two dimensions explain one third of the total variance. Task performance, years of education, as well as the cognitive measures of attention and working memory contribute most strongly to Dimension 1, which we thus termed “Cognition”. By contrast, Dimension 2, is most strongly correlated with the majority of neural measures on the one hand (with negative loadings), and the SSQ questionnaire ratings (with positive loadings), on the other, which we thus termed “Auditory processing”. Critically, participants’ age contributed less than 4 % to these two dimensions, i.e., less than what would be expected if all 25 variables used in the multivariate analysis contributed equally (Fig. 5B).

**Figure 5.**
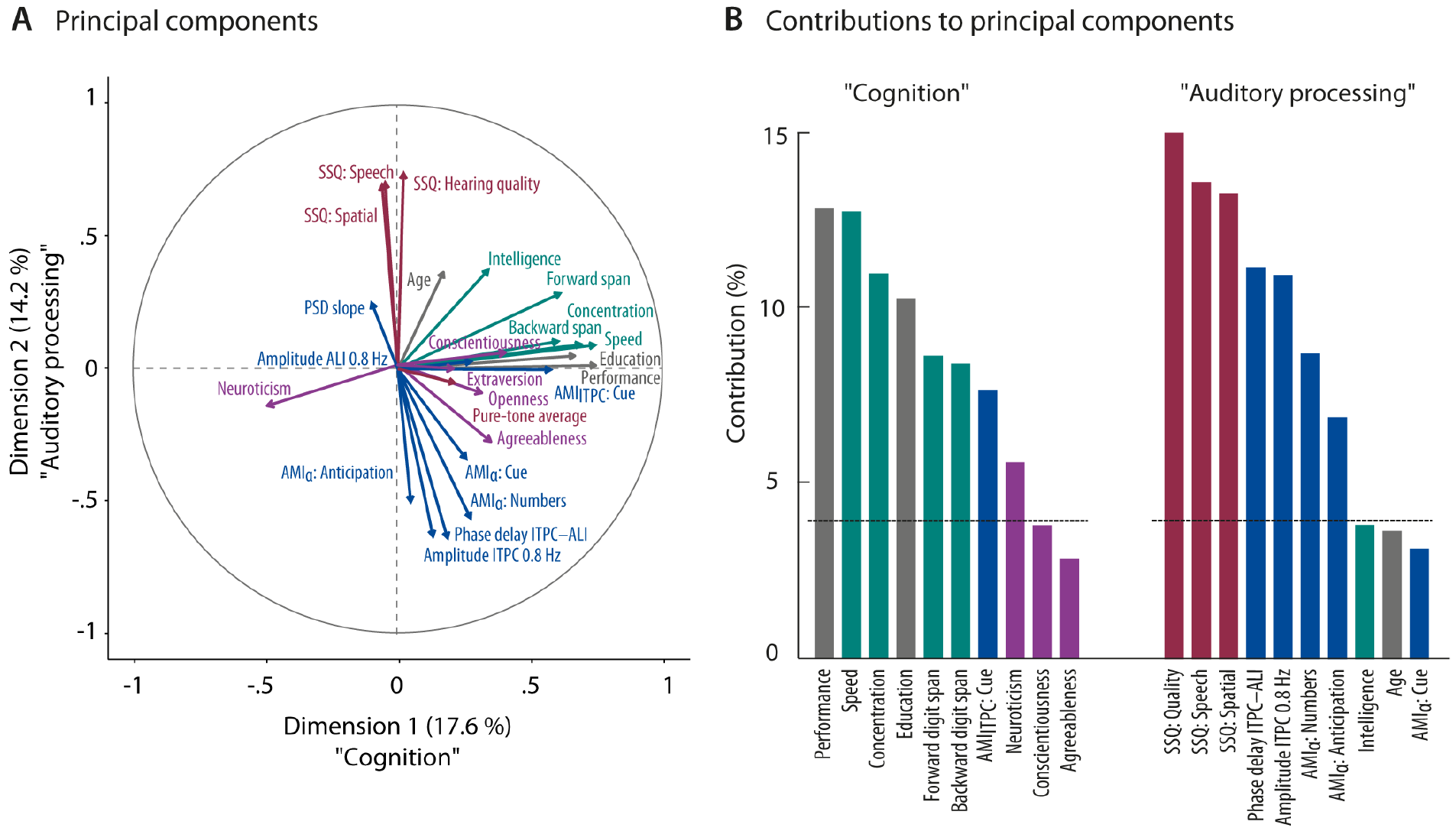
(A) Correlation circle showing the projection of all 25 variables of interest (represented as arrows) into the vector space spanned by the first two principal components. The length and position of an arrow denotes how well the corresponding variable is represented by (i.e. correlated with) the first two principal components. Colors indicate the grouping of variable into the same five broad categories as in (B) (grey = performance and demographics; purple = personality traits; green = cognitive measures; blue = neural measures; maroon = auditory perception). (D) Bar graphs show the ten variables with the highest contribution (in %) to Dimension 1 and 2. Black dotted lines indicate the average contribution (4 %) expected under the assumption of equal contribution of all 25 included variables. Note how variables describing cognitive abilities and neural measures are separated by their contributions to Dimension 1 and 2, respectively. [Color figure can be viewed at wileyonlinelibrary.com]

It is important to note, however, that we focused on the distribution of variables along the first two dimensions because they jointly explain the largest proportion of variance. This does not imply that additional dimensions were not meaningful for the interpretation of multivariate patterns and caution is needed when interpreting the relationships between individual measures based on the projection of variables into this two-dimensional representation, alone.

## Discussion

In this study, we investigated the contribution of neural oscillatory dynamics to the implementation of selective auditory attention with a focus on middle-aged and older adults. Specifically, we asked whether the degree to which participants modulated alpha power in sync with rhythmic auditory input could serve as a neural marker of listening success in the aging individual.

Overall, this sample of middle-aged and older adults (39-69 years) exhibits neural patterns that, to a large degree, mirrored the results previously reported for younger adults (23-34 years; Wöstmann *et al.*, 2016). Similarly, the overall pattern of behavioral results was in line with that observed for younger adults, based on a purely descriptive comparison of the two samples. Possibly due to the slightly faster speech presentation rate chosen here, middle-aged and older adults showed a small decrease in hit rate but still produced significantly more stream confusions than random errors. We found a sustained lateralization of alpha power that was modulated at the frequency of speech presentation and positively related to task performance. Instead of the changes in performance or neural dynamics correlating with age, a strong relationship between performance and an individuals’ attentional and short-term memory capacity emerged. We will now go on to discuss these results in more detail.

### Middle-aged and older adults preserve attentional inhibition and dynamic modulation of lateralized alpha power

Somewhat contrary to our expectations, we found that middle-aged and older adults performed the dichotic listening task almost on par with the younger adults tested in the study by Wöstmann *et al.* (2016). This may appear surprising given the vast amount of evidence pointing to age-related deficits in attentional inhibition (e.g., Hasher & Zacks, 1988; Gazzaley *et al.*, 2005; 2008). However, our results also add to a growing body of evidence suggesting that the ability to spatially direct attention remains relatively unaffected by age (Nissen & Corkin, 1985; Madden, 2007; Mok *et al.*, 2016; for a comprehensive review see Zanto & Gazzaley, 2014). One might claim that the low linguistic load, i.e., the presentation of syntactically and semantically unconnected elements of a closed set of numbers, could have resulted in low distractibility of the irrelevant information. We would argue, however, that the tight temporal alignment of numbers across the two streams that were spoken by the same female talker did in fact lead to overall high levels of energetic and informational masking.

At the neural level, we observed patterns that, to a large extent, replicate the results found for younger adults. That is, we found the hemispheric lateralization of alpha power and its temporal alignment with auditory inputs (here at the presentation rate of 0.8 Hz) to be generally preserved in our sample of middle-aged and older adults. Moreover, we found a significant increase in the 0.8-Hz amplitude of the time-resolved alpha lateralization index (ALI) amplitude for correct vs.incorrect trials. This speaks to a functional role of lateralized alpha dynamics for behavior also in the under-researched population of middle-aged and older adults.

Our findings contrast with previous reports of pronounced decreases in the hemispheric modulation of neural activity in the alpha band with age (Sander *et al.*, 2012; Vaden *et al.*, 2012; Hong *et al.*, 2015; Leenders *et al.*, 2016). There are several possible explanations for the observed discrepancy. As highlighted before, the fact that all of the previous studies concerned with aging effects investigated the role of alpha power modulation in visual working memory renders direct comparisons with the present auditory study difficult. Differences in the compared age groups might be another possible explanation as we purposefully focused on middle-aged and older adults (~40-70 years) to test a broader, potentially more heterogeneous, and overall younger cohort than many of the previous aging studies.

Our results, however, are also to some extent compatible with the findings by Mok *et al.* (2016) and Leenders *et al.* (2016). Mok and colleagues focused explicitly on older adults (i.e., 60-87 years) and observed intact alpha power lateralization and spatial cue benefit for behavior, while Leenders *et al.* (2016) report preserved alpha power lateralization during spatial cueing but not during a subsequent working memory retention interval. Older adults were not found to be more easily distracted by irrelevant information than younger adults. Lastly, the results by Sander *et al.* (2012) revealed an influence of task difficulty on the degree of hemispheric lateralization of alpha band activity with older adults exhibiting the strongest lateralization at an intermediate level of difficulty. Together with the present results, this suggest that selective attention is not generally impaired at older age and that, beyond age, the neural implementation of top-down attentional control hinges on a variety of additional factors.

### Integrity of speech-synchronized alpha power modulations as neural marker of successful listening behavior in the aging adult?

In the present study, we were interested in probing the value of dynamic alpha power lateralization as a predictor of successful adaption to challenging listening situations in the middle-aged and older listener. Specifically, we asked in how far changes in the implementation of spatial attention at the neural level across age would be associated with concomitant changes in behavior.

While we found a significant difference in 0.8-Hz ALI amplitude for correct vs. incorrect trials across all participants, this effect was not modulated by age. Furthermore, this effect relied on a relatively coarse differentiation of fully correct trials and trials with at least one but up to four errors. By contrast, the correlations of a more fine-grained measure of performance, i.e., the proportion of correctly recalled numbers, or age with neural measures describing different aspects of the attentional modulation of alpha oscillations collectively failed to show even a moderate relationship. Instead, performance was best predicted by an individual’s attentional and working memory capacity, attesting to the task-induced demands in attentional control, and to a lesser degree, working memory.

The comparison of results from the aging studies discussed above suggests that the link between the relative degree of lateralized alpha power and associated consequences for behavior across age is indeed an elusive one. That is, the diminished or absent lateralization of alpha power observed for older compared to younger adults is not consistently paired with pronounced differences at the behavioral level (Sander *et al.*, 2012; Hong *et al.*, 2015; Leenders *et al.*, 2016).

Taken together, the results from the present and previous studies suggest that the link between behavior and the fidelity of alpha power lateralization consistently observed for younger adults (Kelly *et al.*, 2006; Thut, 2006; Sauseng *et al.*, 2009; Bengson *et al.*, 2012; Wöstmann *et al.*, 2016) becomes less stable with age. There are several possible explanations for this phenomenon. First of all, as reported by previous studies, alpha dynamics in the broadest sense seem to alter with age, diminishing the likelihood of uncovering nuanced statistical relations. Specifically, it had been shown that older adults are less able to maintain a heightened level of alpha power over longer periods of time (Wöstmann *et al.*, 2015a; Henry *et al.*, 2017). Furthermore, it is possible that the relationship between lateralized alpha power and behavior becomes obfuscated due to the increased utilization of alternative, compensatory processing strategies. Such shifts in processing strategies could be observed at either the cognitive and/or neural level. At the level of cognitive processing strategies, it has been shown that older adults are able to compensate for difficulties in auditory processing by relying more strongly on cognitive strategies and more specifically by capitalizing on top-down processing (Kathleen Pichora-Fuller, 2009; Anderson *et al.*, 2013). At the neural level, these changes in processing mechanisms may be accompanied by the recruitment of additional neural resources such as the increased involvement of prefrontal brain regions to compensate for less efficient processing in low-level sensory areas (Grady, 2012).

In sum, despite our finding of differential modulation of alpha power in correct and incorrect trials, we cannot conclude that the dynamic lateralization of alpha power is a necessary or sufficient neural mechanism supporting selective auditory attention across age. Thus, if we are to understand the mechanisms that govern listening behavior in the aging adult, we cannot restrict our investigations to a study of specific neural measures of interest that becomes detached from concomitant changes at different levels of observation, such as behavior (Krakauer *et al.*, 2017). Instead, we need to adopt a more comprehensive approach that considers the relative contributions of various behavioral and neural factors, and their interactions.

### Changes in cognitive abilities and neural dynamics describe separate dimensions in the aging listener

As highlighted above, if we want to uncover and characterize the neural mechanisms and behavioral strategies that support speech perception and comprehension under unfavorable, i.e., real-life, settings, our investigations must be as broad as possible. Therefore, the present study has adopted a more comprehensive strategy to help understand the functional role of oscillatory dynamics in the aging listener.

Our multivariate analysis of variables including demographic factors, personality traits, cognitive abilities, hearing acuity, as well as task-dependent and -independent neural measures reveal a pattern in which age is a surprisingly poor predictor of task-related neural dynamics or behavior. Instead, the results of the principal component analysis uncovered an underlying structure that separated neural variables and cognitive measures into different dimensions. Again, effects of age were largely orthogonal to these first two dimensions that we termed “Cognition” and “Auditory processing”.

It is well-attested that increased age goes hand in hand with gradual changes at the (neuro–) biological level that are in turn reflected in decline not only in sensory fidelity but also in many aspects of cognitive functions, such as attention, memory and processing speed (Lindenberger & Baltes, 1997; Verhaeghen & Cerella, 2002; Raz, 2005; Grady, 2012). However, it is important to realize that this general trajectory of cognitive decline and its relationship with underlying neural degeneration is accompanied by a considerable amount of inter-individual variability that increases with age (Rapp & Amaral, 1992; Li *et al.*, 2001; Hedden & Gabrieli, 2004; Fabiani, 2012). This observation fits well with the results of the present study as we found a great amount of heterogeneity with respect to cognitive and neural functioning in our sample of middle-aged and older adults that did not allow for any definite conclusion about the link between these levels of observation. Moreover, it highlights the importance of not only recognizing but capitalizing on the inter-individual variability if we want to better understand how neural oscillations support complex higher order cognitive functions—here, spatial attention in service of speech comprehension.

An obvious limitation of the present study lies in its modest sample size. Also, studies like the present one face a potential sampling bias that could have favored selecting highly-functioning older adults; note, however, that in our sample, age was not correlated with the level of education. As such, additional studies that focus on a broader sample of participants (and possibly an even larger age range) will be needed to further validate the current findings.

## Conclusions

We observed top-down spatial attention, indexed at the behavioral level by successful task performance and at the neural level by hemispheric modulation of alpha oscillatory power, to be largely intact in middle-aged and older adults. The absent modulations by age in both behavioral and neural-oscillatory markers of the auditory spatial attention call into question the predictive value of alpha power lateralization for individual listening success in the aging adult. Instead, our results emphasize the fact that the considerable amount of inter-individual variability associated with healthy aging can obscure the relationship between neural and cognitive functions.

## Acknowledgements

This research was supported by an ERC grant (ERC-CoG-2014 AUDADAPT) to Jonas Obleser. The authors are grateful to Franziska Scharata, Philipp Seidel, Elisabeth Ni and Felix Deilmann for their help with data acquisition.

## Author Contributions

S.T, M.W., and J.O. designed research; S.T. performed research; S.T., M.W., and J.O. analyzed data; and S.T., M.W., and J.O. wrote the paper.

## Data Accessibility Statement

Data are available from the corresponding authors upon request.

